# Accelerated lipid catabolism and autophagy are cancer survival mechanisms under inhibited glutaminolysis

**DOI:** 10.1101/230433

**Authors:** Anna Halama, Michal Kulinski, Shaima S. Dib, Shaza B. Zaghlool, Kodappully S. Siveen, Ahmad Iskandarani, Noothan J. Satheesh, Aditya M. Bhagwat, Shahab Uddin, Gabi Kastenmüeller, Olivier Elemento, Steven S. Gross, Karsten Suhre

**Affiliations:** Department of Physiology and Biophysics, Weill Cornell Medicine – Qatar Education City, PO 24144 Doha, Qatar; Translational Research Institute, Academic Health System, Hamad Medical Corporation, PO 3050, Doha, Qatar; Institute of Bioinformatics and Systems Biology, Helmholtz Zentrum München, German Research Center for Environmental Health, 86764, Neuherberg, Germany; Caryl and Israel Englander Institute for Precision Medicine, Institute for Computational Biomedicine, Sandra and Edward Meyer Cancer Center, Weill Cornell Medicine, New York, NY, USA; Department of Pharmacology, Weill Cornell Medicine, Cornell University, New York, NY, USA

**Keywords:** cancer metabolism, glutaminolysis, inhibition of glutaminolysis, beta-oxidation, autophagy

## Abstract

Suppressing glutaminolysis does not always induce cancer cell death in glutamine-dependent tumors because cells may switch to alternative energy sources. To reveal compensatory metabolic pathways, we investigated the metabolome-wide cellular response to inhibited glutaminolysis. We conducted metabolic profiling in the triple-negative breast cancer cell line MB-MDA-231, treated with different dosages of glutaminase inhibitor C.968 at multiple time points. We found that multiple molecules involved in lipid catabolism responded directly to glutamate deficiency as a presumed compensation for energy deficit. Accelerated lipid catabolism, together with oxidative stress induced by glutaminolysis inhibition, triggered autophagy. We therefore simultaneously inhibited glutaminolysis and autophagy, which induced cancer cell death. Our study emphasizes the potential of non-targeted metabolomics to characterize and identify metabolic escape mechanisms contributing to cancer cell survival under treatment. Our findings add to the increasing evidence that combined inhibition of glutaminolysis and autophagy may be effective in glutamine-addicted cancers.

## Introduction

Cancer cells evolve under selective pressure to adjust their metabolism to the high biosynthetic and bioenergetic demands of sustained cell division (Pavlova and Thompson, 2016). The role of cancer-specific metabolic pathways was first identified by Otto Warburg, who reported increased glucose consumption and lactate release in proliferating cancer cells (Warburg et al., 1927). This picture successively expanded with the identification of further cancer-specific metabolic alterations (Pavlova and Thompson, 2016). For instance, some tumors rely on elevated glutaminolysis to cover their energy needs and fill their carbon and nitrogen pools for the biosynthesis of purines, pyrimidines, nonessential amino acids, and fatty acids (Altman et al., 2016). Furthermore, certain lung cancers use branched chain amino acids for protein synthesis and as a source of nitrogen (Mayers et al., 2016). Recently, lipid catabolism, in particular fatty acid oxidation, was recognized as a further central energy-producing pathway in different cancer types, facilitating proliferation, motility, and metastasis (Beloribi-Djefaflia et al., 2016; Caro et al., 2012; Carracedo et al., 2013; Liu et al., 2010; Nath and Chan, 2016; Park et al., 2016). Overall, cancer metabolism appears to be very diverse and driven by intrinsic factors, such as the cancer’s genetic makeup, tissue of origin, stage, and grade, as well as extrinsic factors such as the tumor microenvironment or access to oxygen and nutrients (Vander Heiden and DeBerardinis, 2017). Hence, molecules and processes that contribute to cancer-specific metabolic alterations can serve as targets for therapeutic interventions with the aim of effectively disrupting cancer cell metabolism.

Glutamine metabolism, reported as strongly altered in many cancer types, recently emerged as a promising therapeutic target (Altman et al., 2016). The enzyme glutaminase catalyzes the conversion of glutamine to glutamate, which is then further incorporated into the tricarboxylic acid (TCA) cycle to support the cell’s energetic needs (Moreadith and Lehninger, 1984). In particular, expression of the glutaminase splice variant glutaminase C (GAC) has been associated with a higher glutaminolytic flux in various cancers and indicates glutamine addiction of these cancer cells (Gross et al., 2014; Jacque et al., 2015; Timmerman et al., 2013). Among several compounds targeting glutamine metabolism, inhibitors of glutaminase and GAC, namely CB-839 (Gross et al., 2014) and C.968 (Wang et al., 2010), respectively, are effective in suppressing proliferation of certain breast cancers and hematological malignancies as well as in inhibiting oncogenic transformation (Gross et al., 2014; Stalnecker et al., 2015; Wang et al., 2010). However, cancer cell susceptibility to glutamine inhibition can vary among individual cancers. Indeed, treatment for 72 h of breast cancer cell lines with compound CB-839 induces apoptosis in 60% of HCC1806 cells but in only 20% of MB-MDA-231 cells, although growth rates of both cancer cell lines are significantly affected (Gross et al., 2014). This finding suggests that certain cancer cells can develop survival mechanisms to overcome nutrient insufficiency. In a clinical setting, such mechanisms can lead to the emergence of resistance. Consequently, uncovering the molecular bases of these metabolic cancer survival mechanisms may contribute to the identification of novel targets that allow for combination therapy and can be tailored to the treatment of specific cancer types.

We hypothesized that survival mechanisms that are activated under inhibited glutaminolysis would be associated with generalized changes in metabolic pathways and that these alterations can be captured and characterized by metabolome-wide profiling. We therefore implemented a broad, non-targeted metabolic profiling approach to identify alterations in cancer metabolism that contribute to cancer cell survival under glutaminolysis inhibition. To the best of our knowledge, metabolome-wide profiling has not been previously used to provide insights into the rewiring of cancer cell metabolism in such settings. So far, previous studies monitored only the impact of glutaminase inhibition by measuring a small set of molecules, i.e., glutamate, glutathione, and TCA cycle intermediates. These studies investigated HCC1806 and T47D cell lines treated with CB-839 (Gross et al., 2014) and mouse embryonic fibroblasts treated with C.968 (Stalnecker et al., 2015). Based on the previous studies in which glutaminolysis inhibition in MB-MDA-231 cell line decreased cell proliferation but was insufficient to trigger apoptosis, we suspected metabolic adaptation in the “glutamine-addicted” MB-MDA-231 cells (Gross et al., 2014; Stalnecker et al., 2015). Therefore, we selected the MB-MDA-231 cell line for our study to identify metabolic “survival mechanisms.” For glutaminolysis inhibition, we used a commercially available, allosteric inhibitor of GAC, compound C.968 (Stalnecker et al., 2015; Wang et al., 2010). This compound effectively suppresses MB-MDA-231 cell proliferation, but its effect on apoptosis, necrosis, and perturbation in the cell cycle have not previously been investigated (Stalnecker et al., 2015; Wang et al., 2010). Here, we investigate cellular processes and metabolic alterations in the MB-MDA-231 cell line under treatment with C.968 at two different dosages and at multiple time points (10 h, 24 h, 48 h, and 72 h). We show that inhibition of glutaminolysis alone does not lead to significant levels of apoptosis, necrosis, or cell cycle arrest, which has not been previously investigated (Stalnecker et al., 2015; Wang et al., 2010). Using broad-range metabolic profiling covering molecules from eight primary pathways including the metabolism of amino acids, carbohydrates, lipids, nucleotides, cofactors, vitamins, peptides, and xenobiotics, we identify accelerated lipid catabolism as an event contributing to survival of glutaminase-inhibited cancer cells. Taken together, the observed alterations in metabolic pathways suggested a critical role for autophagy in cell survival, which we subsequently confirmed. Eventually, we show that simultaneous inhibition of glutaminolysis and autophagy increases cell death and thus offers a promising treatment option.

## Results

### Glutaminase Inhibition Reduces Cell Proliferation but Does Not Trigger Apoptosis or Cell Cycle Arrest

In a previous study, C.968 was reported to be a potential glutaminolysis inhibitor in the GAC-expressing triple-negative breast cancer (TNBC) cell line MDA-MB-231 (Katt et al., 2012; Stalnecker et al., 2015; Wang et al., 2010). Our first objective was thus to verify that compound C.968 effectively inhibits glutaminolysis in our experimental set-up. The presence of GAC protein in the MDA-MB-231 cell line was determined using western blot (**Supplementary Figure 1 A**). Glutaminase catalyzes the transformation of glutamine to glutamate, which is then further incorporated into the TCA cycle to support cellular energetic needs (Moreadith and Lehninger, 1984). The inhibition of GAC is expected to reduce the amount of available substrate for oxidation, and a decreased oxygen consumption rate (OCR) under inhibited glutaminolysis thus is expected. We used OCR as the readout for monitoring mitochondrial respiration in the MDA-MB-231 cell line. Toward this end, we quantified OCR in the MDA-MB-231 cells that were treated with either vehicle or 10 μM C.968 after incubation in a conditioned medium supplemented with the following alternative bioenergetics substrates: 1) a combination of glucose plus glutamine; 2) glutamine only; or 3) glucose only. We observed a decrease in OCR in cells treated with C.968 in a medium supplemented with glutamine only, as well as in glucose-plus-glutamine medium, but not in medium supplemented with glucose (Figure 1A). The observation of higher OCR in the untreated cells in medium containing glutamine vs. glucose indicates that MDA-MB-231 cells rely on glutamine as their principal mitochondrial fuel source, in agreement with a previous report suggesting the addiction of TNBC to glutamine (Gross et al., 2014). A greater OCR decrease in TNBC cells cultivated in media with glutamine further indicated that C.968 inhibited GAC.

**Figure 1.**
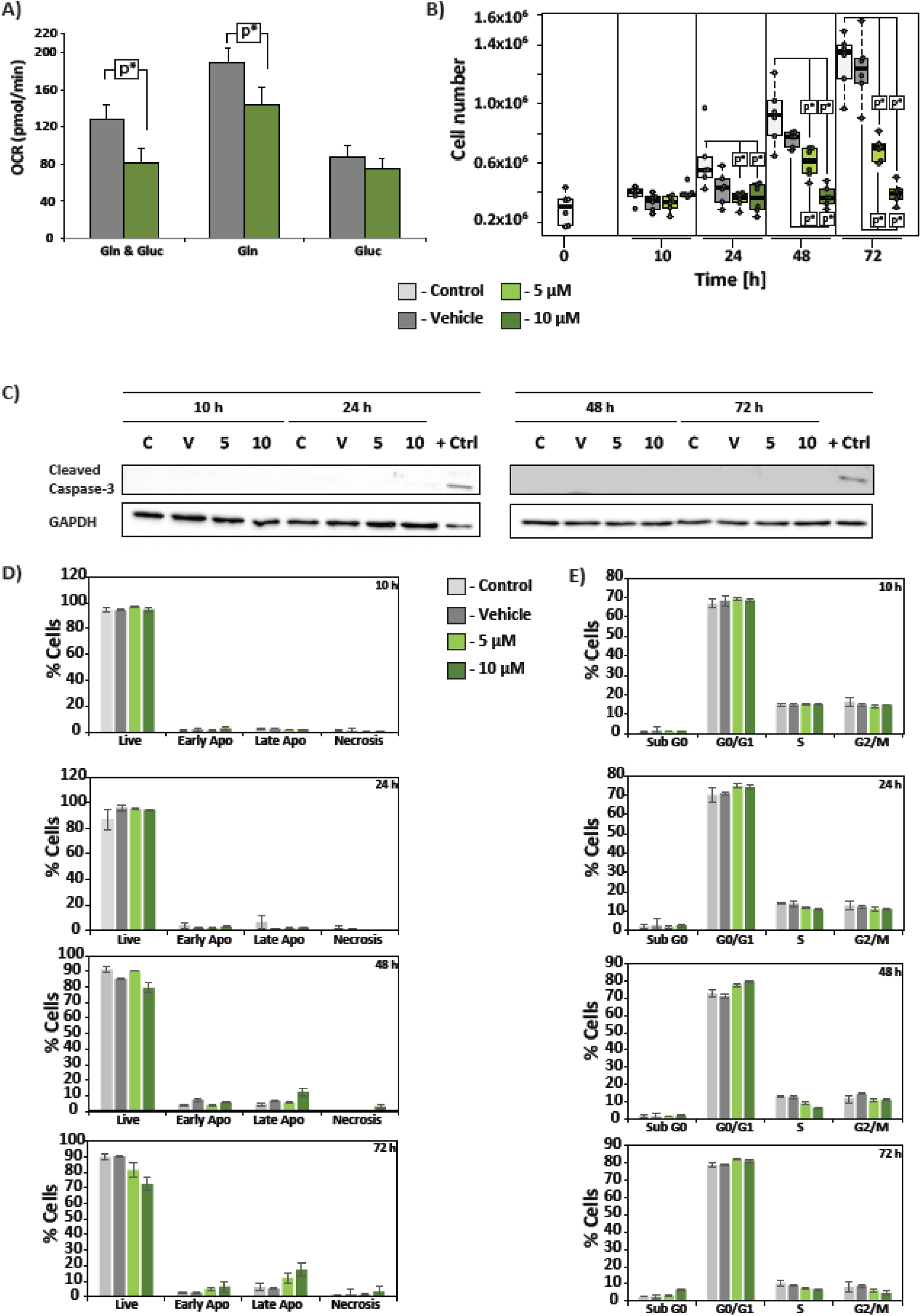
C.968 inhibits proliferation of MDA-MB231 cells but does not induce apoptosis or cell cycle arrest. **A)** Oxygen consumption rate (OCR) was determined in cells treated with vehicle or 10 μM C.968 after cell incubation in media containing: 1) mixture of glutamine and glucose (Gln & Gluc); 2) glutamine only (Gln); or 3) glucose only (Gluc). **B)** Cell number was determined in untreated cells or cells treated with vehicle or C.968 (5 μM and 10 μM) over 72 h. **C)** Western blotting analysis for cleaved caspase-3. **D)** Apoptosis and necrosis were not induced by treatment with C.968 as determined after staining of cells with fluorescein-conjugated annexin-V and propidium iodide (PI) and analysis with flow cytometry. **E)** No changes in cell cycle were observed after treatment, as shown by PI staining and analysis for DNA content by flow cytometry. *p < 0.01; untreated cells are depicted in white, vehicle-treated in grey, and treated with 5 μM or 10 μM C.968 in green and dark green, respectively.

Previous studies have also shown that C.968 effectively suppresses proliferation of MDA-MB-231 cells (Wang et al., 2010). Thus, we monitored the impact of C.968 on MDA-MB-231 cell proliferation over a period of 72 h at multiple time points (10 h, 24 h, 48 h, and 72 h). Treatment with C.968 resulted in a time-and dose-dependent decrease in cell number (Figure 1B), in agreement with previous findings (Wang et al., 2010). A significant decrease in cell number together with morphological changes, namely from a fibroblast-like structure to spherical, was further observed by microscopy investigation, with the largest effect after 72 h of treatment (**Supplementary Figure 1 B**).

We next asked whether the observed decrease in MDA-MB-231 cell number after treatment with C.968 is due to cell apoptosis or necrosis because another glutaminase inhibitor, CB-839, was reported to induce an apoptotic response in only 20% of an MDA-MB-231 cell population (Gross et al., 2014). Accordingly, we quantified apoptosis in the MDA-MB-231 cells over a period of 72 h at different time points by monitoring both the caspase 3 cleavage and Annexin V/propidium iodide (PI) cell membrane dual staining. Neither cleavage of caspase 3 nor Annexin V/PI dual staining was observed within the range of C.968 concentrations used, suggesting no apoptosis in glutaminase-inhibited MB-MDA-231 cells over a 72-h period (Figure 1C, Figure 1D). In agreement with a previous study (Gross et al., 2014), we observed late apoptosis in less than 20% of cells at 48 h and 72 h after C.968 treatment (Figure 1D). Using DAPI staining, we further confirmed the absence of apoptosis after treatment (**Supplementary Figure 2**), using Annexin V/PI dual staining we excluded necrosis (Figure 1D).

As decreased cell proliferation could result from perturbations of the cell cycle, we investigated whether the observed decrease in cell number was associated with cell cycle arrest. We characterized the cell cycle of C.968-treated MB-MDA-231 cells and observed no differences in the cell cycle between treated and untreated cells at 10 h and 24 h after treatment (Figure 1E), when the majority of the cell population was in the G0/G1 phase. The population of cells in the sub-G0 phase was found to increase 72 h after treatment (from 2.42% in vehicle-treated cells to 6.68% in cells treated with 10 μM C.968), with a simultaneous decrease in G2/M phase (from 8.85% in vehicle-treated cells to 4.78% in in cells treated with 10 μM C.968). Taken together, we showed that treatment with C.968 suppresses proliferation of MB-MDA-231 cells but does not induce apoptosis or lead to substantial cell cycle arrest.

### Signatures of Metabolic Adjustment of MB-MDA-231 Cells after Glutaminolysis Inhibition

Because C.968 treatment reduced proliferation, but was insufficient to induce cell death, we hypothesized that inhibition of glutaminolysis in MB-MDA-231 cells triggered a metabolic compensation mechanism that enabled the cells to survive under substrate-limiting conditions (Figure 1). We therefore conducted non-targeted metabolic profiling to obtain a readout of metabolic changes associated with cell responses to C.968 treatment. Using the Metabolon platform (Evans et al., 2009), we quantified relative levels of 346 distinct metabolites for untreated MB-MDA-231 cells as well as those treated with dimethyl sulfoxide (DMSO) (vehicle) and with C.968 at 5 μM and 10 μM for time points of 10 h, 24 h, 48 h and 72 h. The identified metabolites cover molecules from eight primary pathways related to the transformation of amino acids, carbohydrates, cofactors and vitamins, energy, lipids, nucleotides, peptides, and xenobiotics (**Supplemental Table 1**). Treatment of MB-MDA-231 cells with C.968 resulted in time-and dose-dependent alterations in cell metabolism, as revealed by orthogonal partial least squares (OPLS) regression modeling (Figure 2A). The biological replicates clustered tightly together and separated by dosage and duration of treatment. Untreated and vehicle-treated cells grouped together, suggesting only minor effects of DMSO at vehicle concentration (**Supplementary Table 2**). A clear separation of cells treated with 5 μM and 10 μM C.968 and location of the 5 μM cluster between the DMSO and the 10 μM clusters indicated a dose-dependent impact of glutaminase inhibition on the global metabolic profiles. A distinct separation between the time points and their chronological order indicated that progressive metabolic changes induced by glutaminase inhibition took place over time. The corresponding loading plots allowed for identification of those metabolites that contributed most to the separation of these groups, i.e., metabolites aligned with the dosage direction and with the temporal direction in these plots are those of interest for the respective factor (Figure 2B). The loading plot thus suggests that molecules involved in metabolism of lipids, amino acids, and nucleotides were the major contributors to this separation of dosage clusters.

**Figure 2.**
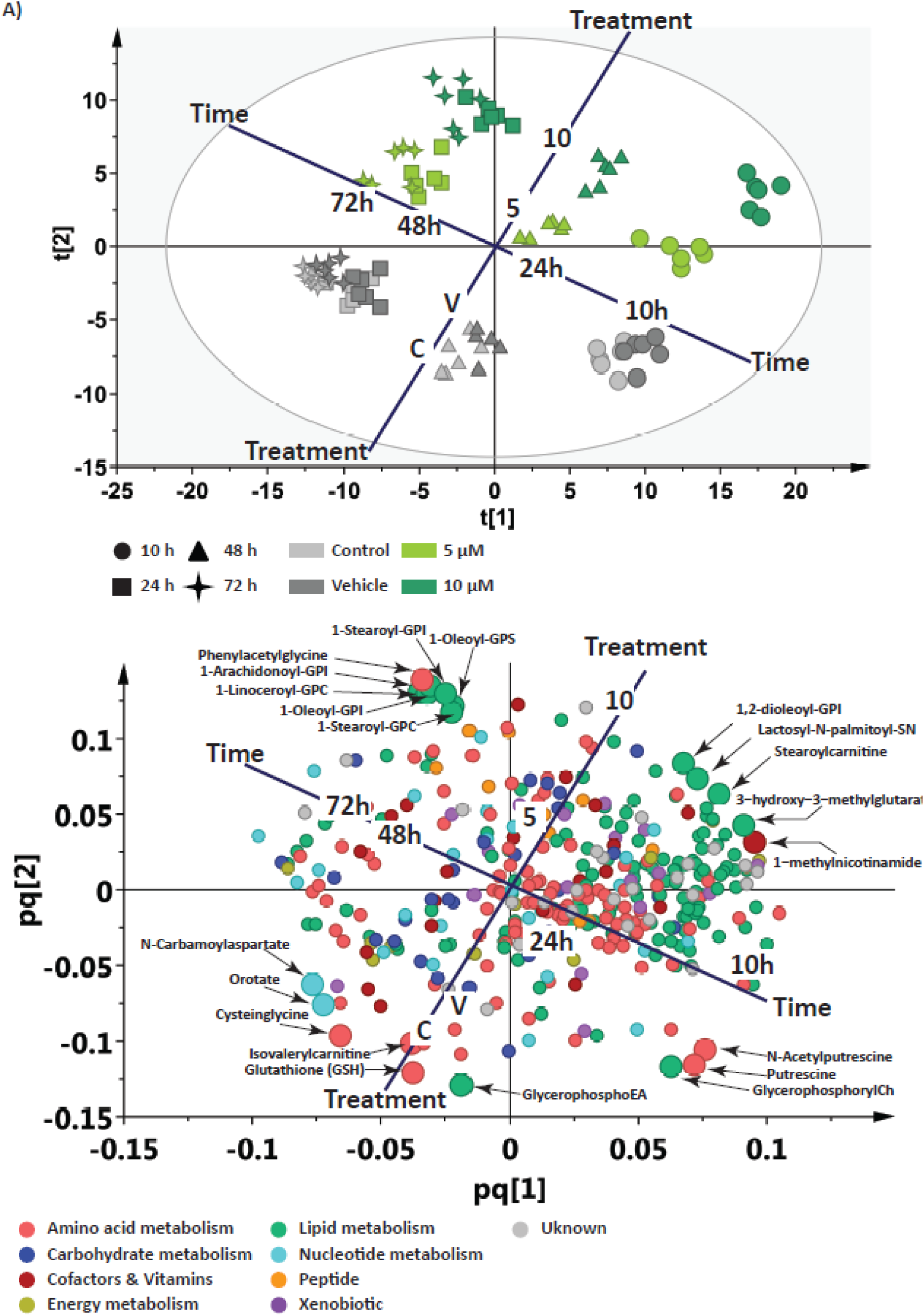
Inhibition of glutaminolysis with C.968 causes general metabolic adjustments. **A)** OPLS reveals time-and dose-dependent separation between groups. **B)** OPLS loading plot shows contribution of different metabolic classes to the separation.

To uncover metabolic alterations that are significant after treatment of the MB-MDA-231 cells with C.968, we applied a linear regression model between the metabolite levels and the dosage of C.968, using time as a covariate. We identified 82 metabolite associations that exhibit a stringent Bonferroni-corrected p-value (p ≤ 1.44 × 10^−4^, corresponding to an alpha error significance level of p ≤ 0.05 after accounting for 346 tests) (Table 1). Most molecules with significantly changed levels were involved in the metabolism of lipids (38 molecules) and amino acids (21 molecules).

**Table 1.**
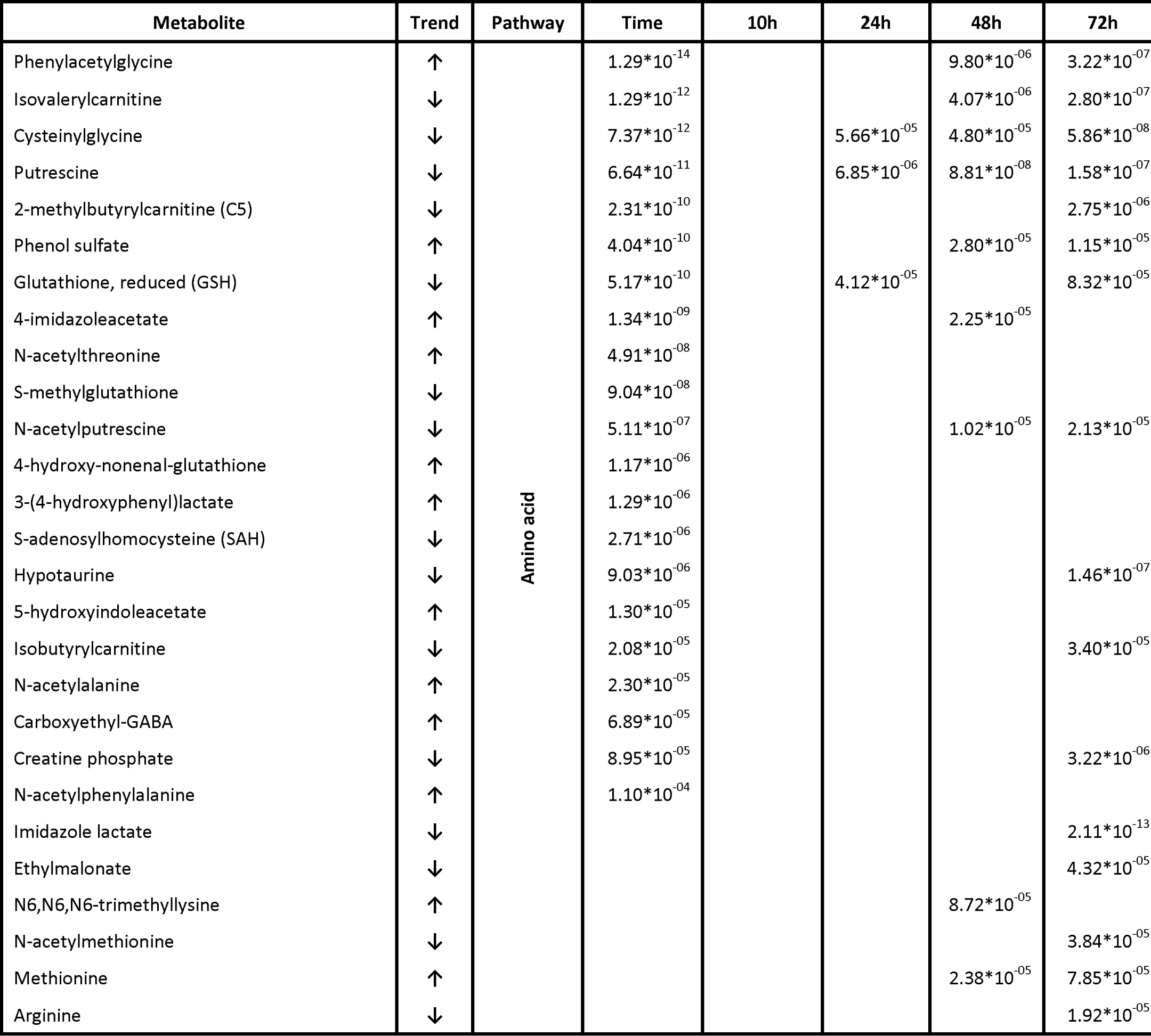
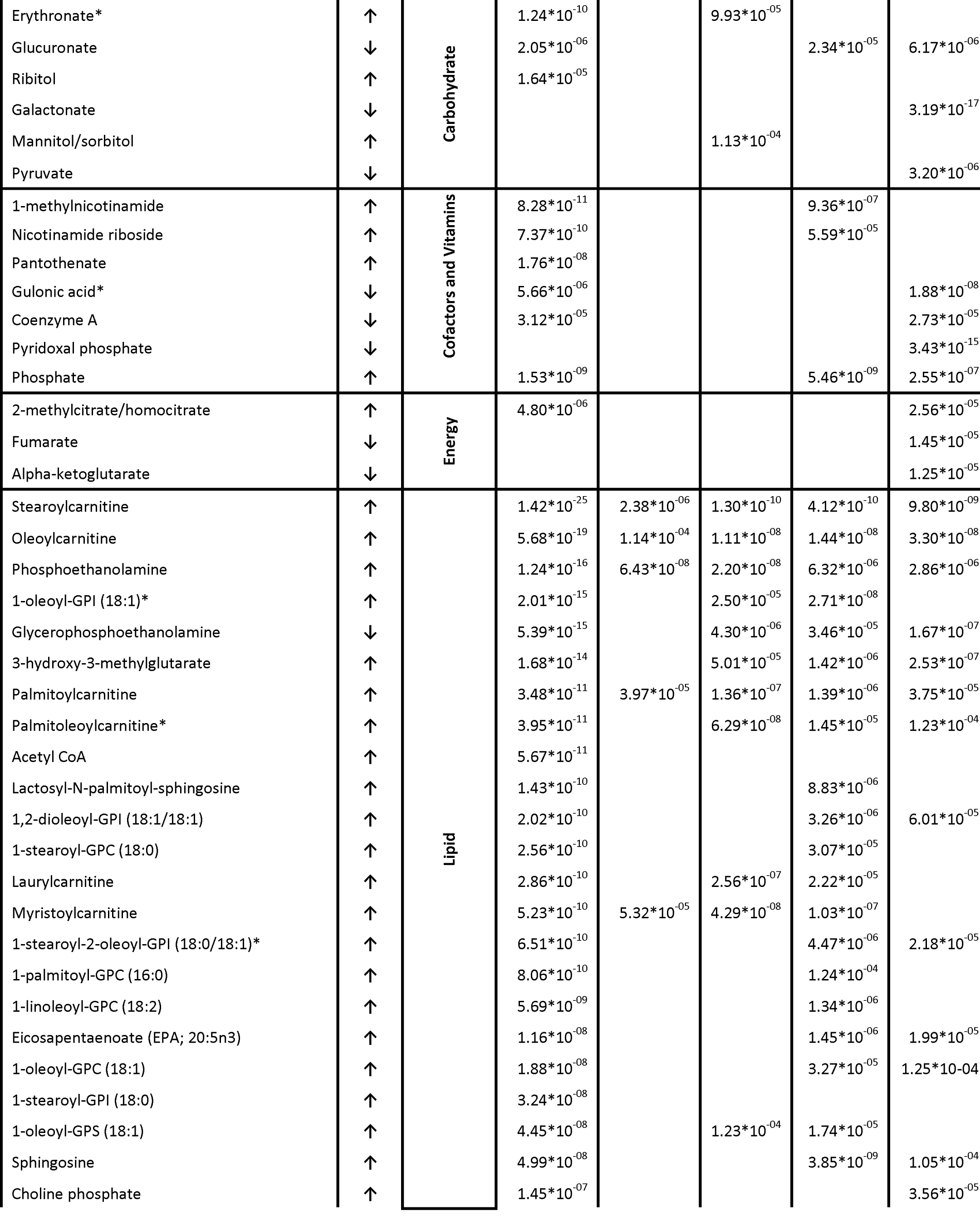
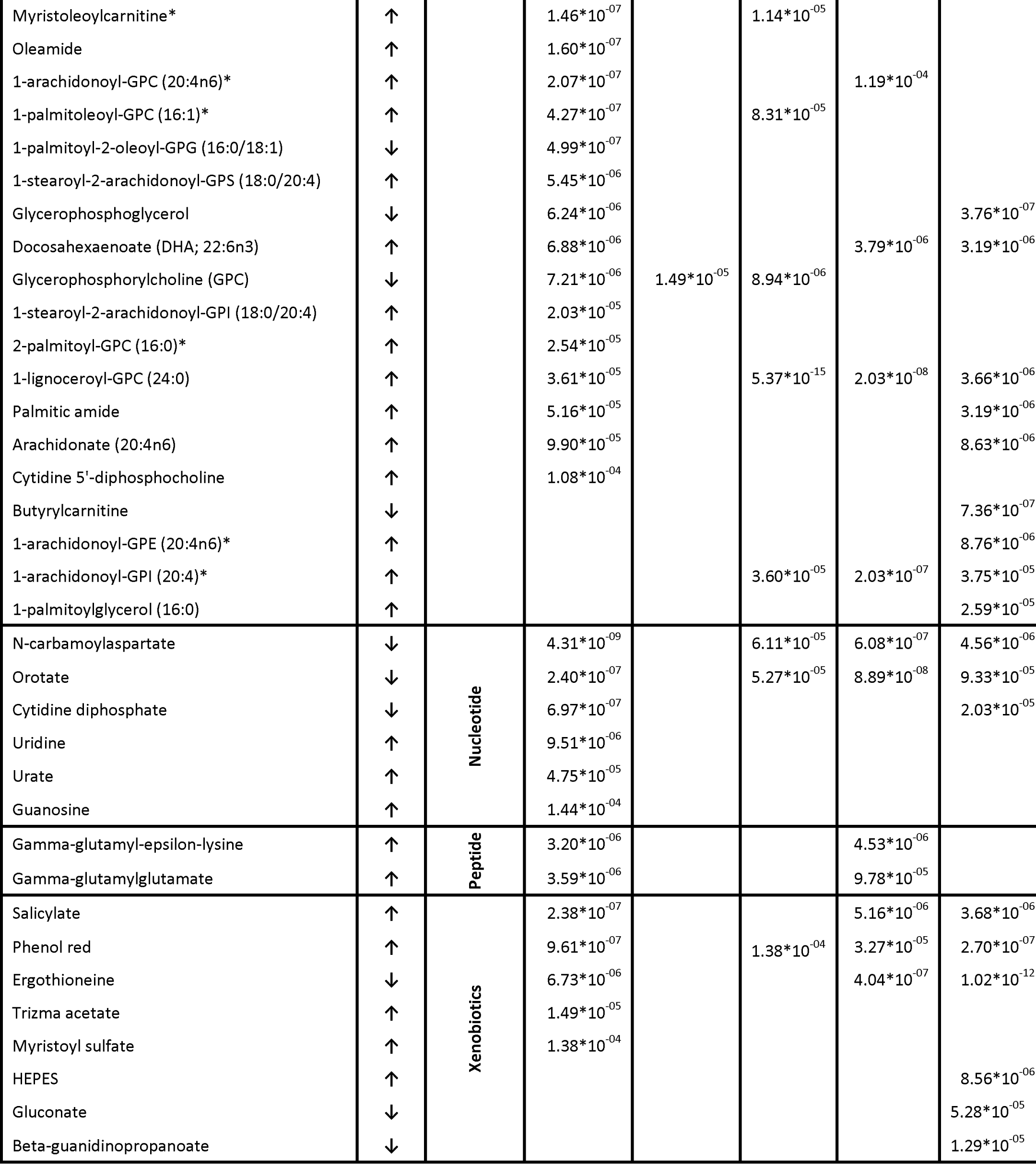
Metabolites altered in C.968-treated MDA-MB-231 cells. The direction of metabolic alterations is represented by arrows (↓ - decrease; ↑ - increase). “Time” and time intervals (10 h, 24 h, 48 h, and 72 h) reflect metabolites showing significant alteration over the entire experimental period and at specific time points (relative to vehicle), respectively. The p-vale was calculated from a linear regression model between the metabolite and dose of C.968, with time as a covariate. The p-value reflects changes in significance observed after cell treatment with 10 μM drug concentration over time.

The metabolic response of MB-MDA-231 cells to C.968 treatment progressed over time. During the first 10 h after treatment, only 6 metabolites were significantly altered, all lipid metabolism intermediates. After 24 h of C.968 treatment, the number of significantly altered metabolites increased to 24 molecules, involved in the metabolism of lipids, amino acids, carbohydrates, or nucleotides. We observed significant alterations in 44 and 58 metabolites after 48 h and 72 h of treatment, respectively. Although the altered metabolites were from different metabolic classes at these later time points, the predominately observed changes were with molecules involved in lipid and amino acid metabolism. These findings suggest that inhibition of glutaminolysis predominantly affects lipid and amino acid metabolism, indicating a central role for these molecules in the metabolic adjustment to glutamate inhibition.

### TCA Cycle Intermediates, But Not Glutamate, Decrease with Inhibition of Glutaminolysis

Inhibition of GAC would be expected to cause a decrease in glutamate level if this pathway was the single cellular source of glutamate. In a previous study, inhibition of GAC with C.968 in mouse embryonic fibroblasts resulted however in no significant depletion of glutamate and TCA cycle intermediate levels, although a significant decrease in ^13^C isotopic enrichment from [U-^13^C]-glutamine was observed for these metabolites (Stalnecker et al., 2015). In agreement with these observations, we found a slight increase in glutamine levels (Figure 3 A) with C.968 treatment, but no significant changes in glutamate levels (Figure 3 B). However, we observed a significant decrease in glutathione (Figure 3 C) and TCA cycle intermediates at 72 h after treatment, including alpha-ketoglutarate, the direct metabolic product of glutamate, along with fumarate and malate (Figure 3 D, E, and F). Other TCA cycle intermediates (citrate, isocitrate, and succinate) were unchanged by GAC inhibition (Figure 3 G, H, and I). A decrease in TCA cycle intermediates without significant changes in glutamate level suggests that glutamate metabolism, rather than absolute glutamate levels, was altered by GAC inhibition in the MB-MDA-231 cell line. These observations indicate that alternative metabolic pathways were activated to compensate for the loss of glutamate and consequent TCA cycle intermediates depletion.

**Figure 3.**
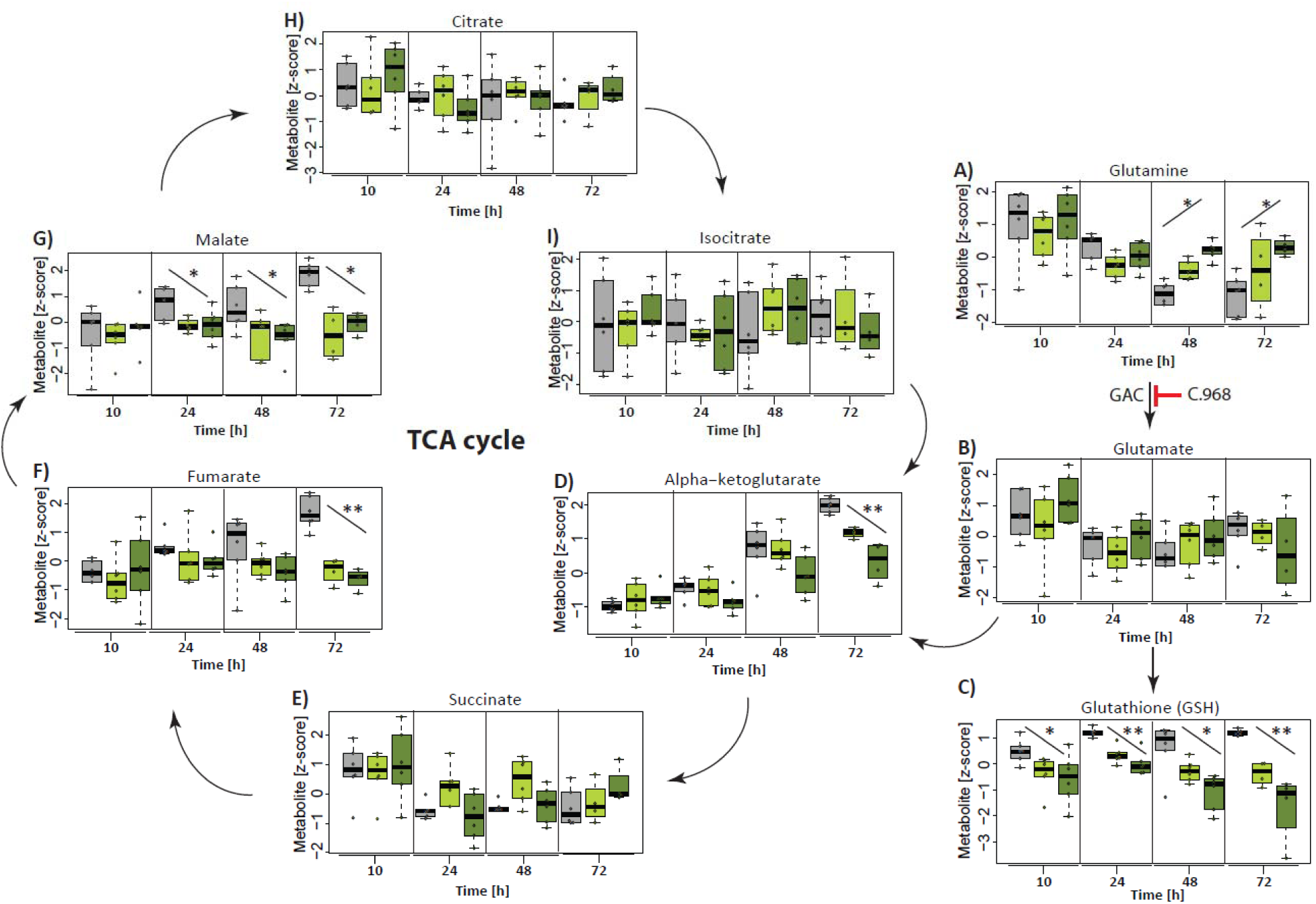
Impact of glutaminase inhibition on the glutamate pool and TCA cycle metabolism. The boxplots present alteration patterns of glutamine, glutamate, reduced glutathione, and TCA cycle intermediates after treatment of cells with C.968. Vehicle-treated cells are depicted in grey, and those treated with 5 μM or 10 μM C.968 are indicated by green and dark green, respectively. ** indicates Bonferroni-corrected significant (p-value ≤ 1.35 × 10^−04^) changes; * indicates trend (p-value ≤ 0.05).

### Lipid Catabolism Accelerates as a Response to Glutamate Depletion

As we have previously shown, associations with ratios between metabolite levels can reveal biochemical pathways and reduce variance within the experimental data (Petersen et al., 2012). To identify metabolites directly affected by treatment of MB-MDA-231 cells with C.968, we therefore calculated metabolic ratios of glutamate relative to all other quantified metabolites. This analysis revealed significant increases (p-value ≤ 8.37⍰×⍰10^−7^; p-gain⍰≥⍰5.96⍰×⍰10^5^) in the ratios of glutamate vs. stearoylcarnitine, oleoylcarnitine, 1-oleoyl-GPI (18:1), and 1-methylnicotinamide (Figure 4). Of note, all four of these metabolites contribute to lipid metabolism: acyl-carnitines are modified fatty acids for traffic across the inner mitochondrial membrane; glycophosphoinositols are lipid species involved in cell structure and signaling processes; and 1-methylnicotinamide is a product of the metabolism of NAD/NADP that contributes to *de novo* lipid biosynthesis and cellular antioxidant defenses. These results suggest the potential involvement of accelerated lipid metabolism as a compensatory response of cancer cells after GAC inhibition.

**Figure 4.**
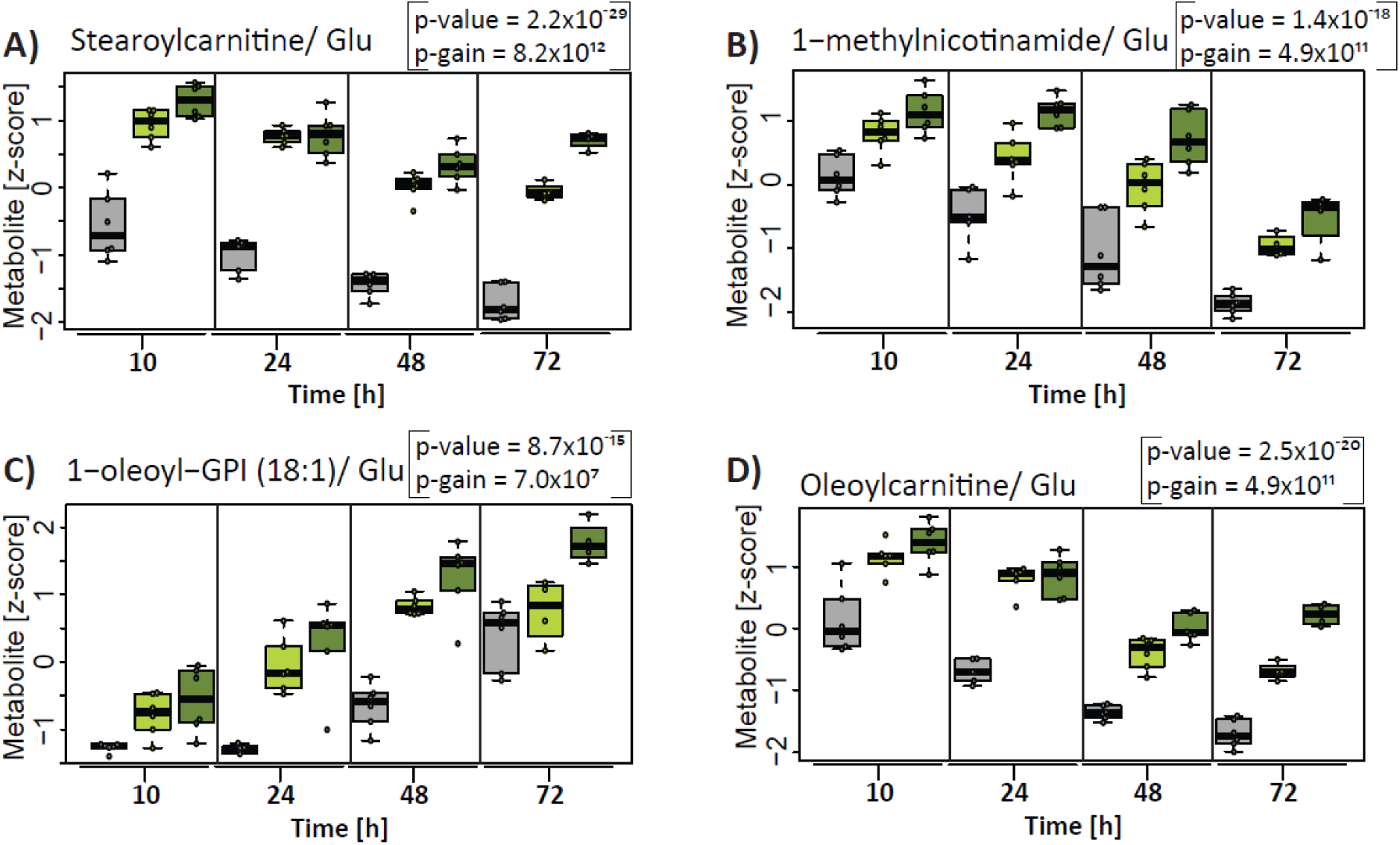
Inhibition of glutaminolysis triggers an increase in ratios between molecules involved in lipid metabolism and glutamate. The boxplot presents significant alterations in the metabolic ratios of glutamate, which were observed over the entire experimental period after cell treatment with C.968. Vehicle-treated cells are depicted in grey, and those treated with 5 μM or 10 μM C.968 are indicated by green and dark green, respectively.

Thus, we further focused on molecules involved in lipid metabolism, showing significant alterations after glutaminolysis inhibition in the MB-MDA-231 cell line. Of note, GAC inhibition resulted in increased levels of lysophospholipids and polyunsaturated fatty acids (PUFA), concomitant with decreases in glycerophosphorylcholine (GPC) and phosphoethanolamines (**Supplementary Figure 3**). These alterations suggest accelerated catabolism of glycerophospholipids. Furthermore, the observed increase in monoacylglycerols suggests accelerated degradation of triacylglycerols and/or diacylglycerols, resulting in the release of fatty acids for ATP generation via beta-oxidation. In accord with this possibility, we observed significant increases in acyl-carnitines, known to deliver fatty acids to the mitochondria for beta-oxidation (**Supplementary Figure 3**). Also, levels of stearoyl-carnitine, palmitoyl-carnitine, myristoyl-carnitine, and oleoyl-carnitine all significantly increased at 10 h after C.968 treatment, followed by significant increases in palmitoleoyl-carnitine, lauryl-carnitine, and myristoleoyl-carnitine at later times (**Supplementary Figure 3**, summarized in Figure 5A).

**Figure 5.**
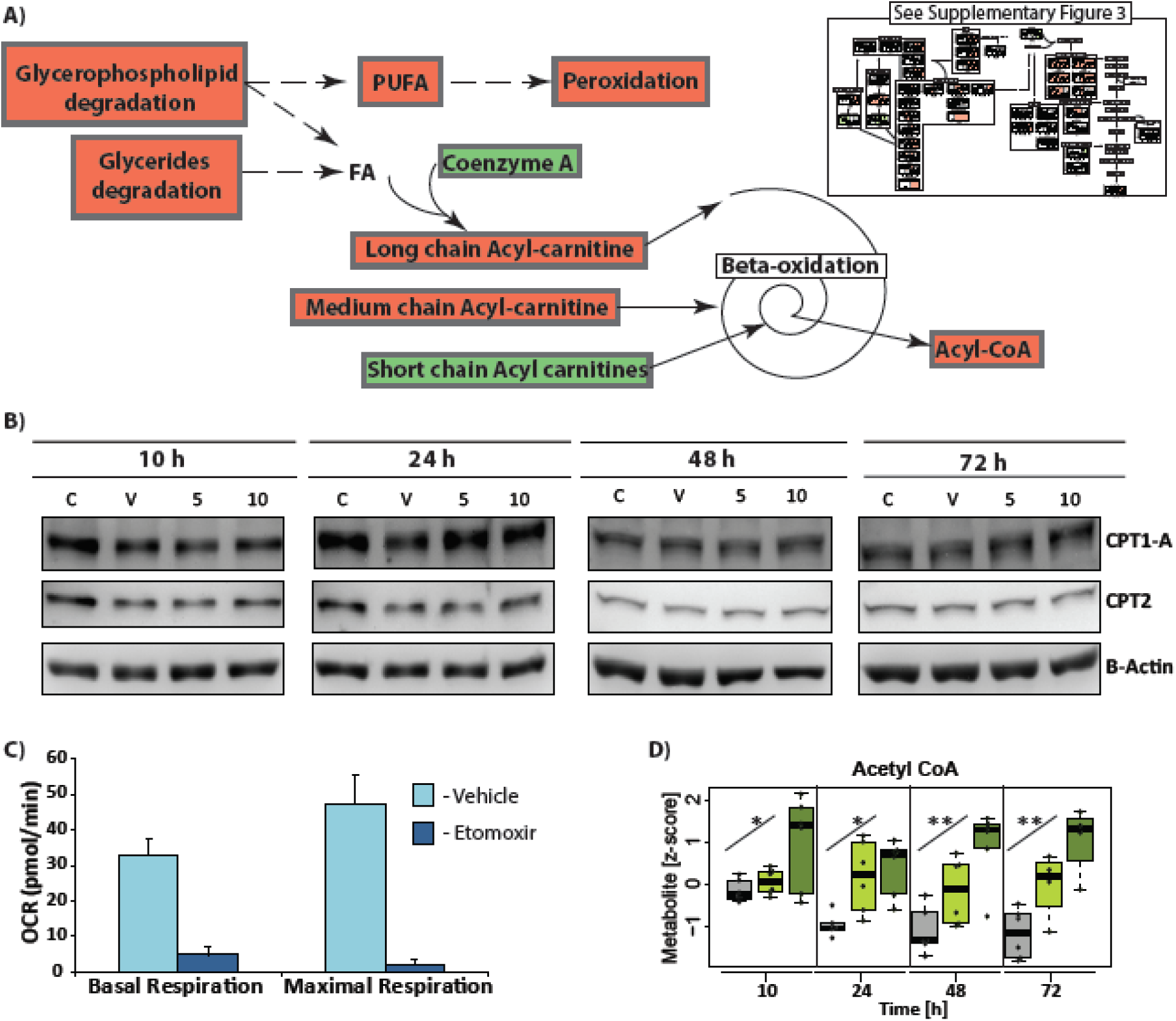
Lipid catabolism is accelerated after glutaminase inhibition. **A)**Overview of lipid metabolism alterations after treatment of MB-MDA-231 cells with C.968. **B)** Western blot of CPT1A and CPT2. **C)** Basal and maximal OCR were monitored in untreated cells and cells treated with CPT1 inhibitor [etomoxir (ETO)] after cell incubation in medium supplemented with bio-available palmitate. **D)** Boxplot showing significant increase in acetyl-CoA, a final product of beta-oxidation. Light blue, vehicle-treated cells at basal oxidation; dark blue, etomoxir-treated cells; grey, vehicle-treated; green, 5 μM C.968– treated cells; dark green, 10 μM C.968–treated cells. *, trend to increase (p-value ≤ 0.05) at the specified time point; **, Bonferroni-corrected significant alteration (p-value ≤ 1.35 × 10^−04^) at the specified time point.

Accumulation of acyl-carnitines carrying long-chain fatty acids can result from either an increase in beta-oxidation or a deficiency in carnitine palmitoyltransferase (CPT) II, the enzyme that catalyzes removal of the carnitine and simultaneous coupling of coenzyme A, the triggering step for beta-oxidation of fatty acids in mitochondria. To address the latter possibility, we assessed the relative abundance of both CPT1A and CPT2 by western blot analysis over a period of 72 h after cell treatment with C.968 (quantified at 10 h, 24 h, 48 h, and 72 h). GAC inhibition did not result in any detectable change in levels of CPT1A or CPT2 (Figure 5B), indicating that beta-oxidation was accelerated in the MB-MDA-231 cells as a response to C.968 treatment.

We next sought to quantify the extent to which MB-MDA-231 cells use fatty acids as an energy source. To test this, we performed a fatty acid oxidation (FAO) assay using a Seahorse fluxometer (Agilent Technology). Toward this end, mitochondrial OCR was measured as a readout of FAO after incubation of cells in media supplemented with bio-available palmitate as the sole bioenergetics fuel source. To confirm that the OCR quantified in this study was specially used for FAO, we inhibited CPT-1, the rate-limiting enzyme required for FAO, with the selective inhibitor etomoxir (ETO). A higher level of basal and maximal OCR in untreated cells vs. ETO-treated cells confirmed that MB-MDA-231 cell could use FAs as a source of energy (Figure 5C). Moreover, we observed a significant increase in cellular acetyl-CoA levels, the final product of beta-oxidation, over the experimental period (Figure 5D). Collectively, these results demonstrate that lipid catabolism is accelerated in response to GAC inhibition, further suggesting that under substrate-limiting conditions, lipid catabolism is accelerated to compensate for loss of energy and to contribute to the TCA cycle in the form of acetyl-CoA.

### Autophagy Is Triggered to Protect Cells from Death after Glutamate Shortage

Lipid catabolism was previously recognized as an important alternative source of energy for cancer cells (Carracedo et al., 2013) and has been associated with autophagy activation (Singh et al., 2009; Wen et al., 2017). A role for autophagy in protecting cancer cells from death under oxidative stress, as well as nutrition insufficiency, is well known (Levy et al., 2017). Because we observed decreased levels of the critical cellular antioxidant glutathione (GSH) following cell treatment with C.968, this result suggests GAC inhibition–mediated oxidative stress (Figure 3 A). In support of this possibility, we showed a significantly diminished ratio of reduced glutathione (GSH to oxidized glutathione) (GSSG), providing a direct readout of C.968 treatment–induced oxidative stress (Figure 6A). Some products of oxidative stress are highly reactive and can chemically modify the cell membrane bilayer by causing peroxidation of polyunsaturated fatty acids (PUFA), leading to formation of lipid hydroperoxides such as 4-hydroxynonenal (4-HNE), as primary products (Ayala et al., 2014). In accordance with GAC inhibition– induced oxidative stress, we observed an increase in 4-hydroxy-nonenal-glutathione, the glutathione conjugate of 4-HNE (see Figure 6B). Together, these findings strongly indicate that MB-MDA-231 cells experience oxidative stress in response to GAC inhibition.

**Figure 6.**
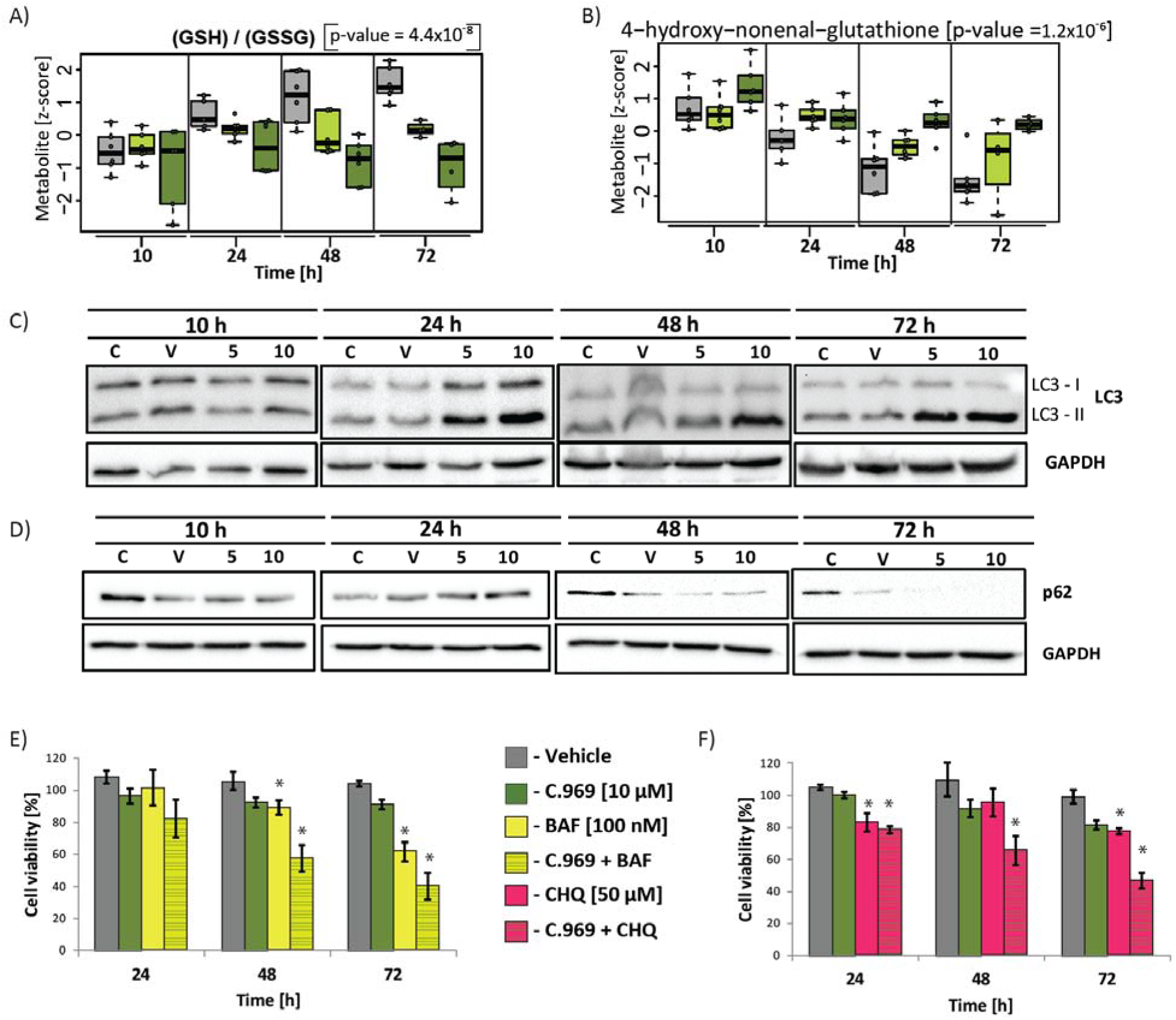
Inhibition of GAC and autophagy triggers cancer cell death. **A)** Boxplot showing significant decrease in ratio between glutathione reduced (GSH) and glutathione oxidized (GSSG), which was observed over the entire experimental period. The p-value reflects significant changes observed after cell treatment with 10 μM drug concentration over time. **B)** Boxplots showing alteration in 4-hydroxy-noneal-glutathione, byproduct of reactive oxygen species targeting n-6 fatty acid. **C)** Western blot for LC3-I and LC3-II after treatment of MB-MDA-231 cells with C.968 at different time points. **D)** Western blot for p62 after treatment of MB-MDA-231 cells with C.968 at different time points. **E, F)** Cell viability determined with MTT assay. Grey, vehicle-treated cells; green, cells treated with 10 μM C.968; yellow, cells treated with bafilomycin (100 nM); greenish-yellow, cells treated with 10 μM C.968 and bafilomycin (100 nM); pink, cells treated with chloroquine (100 μM); greenish-pink, cells treated with 10 μM C.968 and chloroquine (100 μM). Significant changes between control and autophagy-treated cells and control and cells treated with GAC and autophagy inhibitors is indicated by *.

The observation of accelerated lipid catabolism, together with oxidative stress that is not associated with apoptosis or cell cycle aberrations, suggested that cell autophagy may be activated as a driver of lipid catabolism and thus bioenergetics fuel, protecting GAC-inhibited cancer cells from death. During autophagy, the cytosolic form of microtubule-associated protein 1A/1B-light chain 3 (LC3-I) is converted to LC3-II, providing a molecular marker for monitoring the extent of autophagy activation (Kabeya et al., 2000). Our results reveal an increase in LC3-II 24 h after GAC inhibitor treatment, with prominent effects at 48 h and 72 h post-treatment (Figure 6C). Activation of the autophagy machinery is also associated with decline in p62 (sequestosome-1) protein levels, selectively degraded by autophagy (Bjorkoy et al., 2005). Consistent with glutaminase inhibition-induced autophagy, we observed a decrease in the level of p62 in MB-MDA-231 cells after treatment with C.968 for 48 h and 72 h (Figure 6D). These results demonstrate that C.968 treatment induces autophagy in MB-MDA-231 cells.

### Simultaneous Inhibition of Glutaminolysis and Autophagy Increases Cancer Cell Death

We next considered the possibility that simultaneous inhibition of GAC and autophagy would increase cancer cell death. To test this hypothesis, we studied the actions of two potent autophagy inhibitors, chloroquine and bafilomycin A1 (Yang et al., 2013). Chloroquine is an US Food and Drug Administration approved drug for treatment of malaria and inhibits autophagic degradation in the lysosomes (Homewood et al., 1972), whereas bafilomycin inhibits autophagy by blocking fusion between autophagosome and lysosome (Yamamoto et al., 1998). The viability of MB-MDA-231 cells was monitored after simultaneous inhibition of GAC with 10 μM C.968 in combination with chloroquine (50 nM) (Figure 6E) or bafilomycin (100 μM) (Figure 6F). We observed that combined GAC/autophagy inhibitor treatment of MB-MDA-231 results in a significantly greater decrease in cell viability than observed with any single agent. The strongest decrease in cell viability was observed at 72 h after treatment. These findings demonstrate that combination therapy with inhibitors of both GAC and autophagy can substantially reduce cancer cell viability.

## Discussion

Addiction of cancer cells to glutamine and increased glutaminolysis have been recognized as promising targets for selective cancer treatment (Altman et al., 2016). This possibility is supported by the effective anti-proliferative effect of GAC inhibition by C.968 on cancer cells, with little or no effect on normal cells (Stalnecker et al., 2015; Wang et al., 2010), and has been confirmed in a study with a different glutaminolysis inhibitor (CB-839) that recently entered a phase I clinical trial for cancer therapy (Garber, 2016). However, inactivation of a single metabolic pathway in a highly complex metabolic network may be insufficient to trigger cancer cell death due to metabolic compensations, as observed for other monotherapies (Lopez and Banerji, 2017). In this study, we showed that C.968 can suppress the proliferation of MB-MDA-231 cells, in agreement with a previous report (Wang et al., 2010); however, this treatment failed to induce apoptotic or necrotic cell death over the experimental period. Thus, we hypothesized that compensatory metabolic alterations in response to glutamate shortage may contribute critically to cell survival.

Here we demonstrated that metabolome-wide profiling, applied to monitor metabolic consequences of inhibited glutaminolysis, led us to identify compensatory metabolic pathways that contribute to cancer cell survival. Our observation of accelerated lipid catabolism, as a major energy-providing pathway activated under glutaminolysis inhibition, agrees with previous studies showing that under conditions of metabolic stresses (e.g. hypoxia or nutrition shortage) cancer cells can retain viability by oxidizing lipids (Kamphorst et al., 2013; Young et al., 2013). Moreover, inhibition of lipolysis has been shown to sensitize resistant lung adenocarcinoma to chemotherapy (Li et al., 2013), suggesting that fatty acid oxidation is indeed a metabolic pathway that contributes to cancer cell resistance (Carracedo et al., 2013). Our metabolic profiling provides further evidence that lipid catabolism, triggered in response to inhibited glutaminolysis, is a potential cancer “survival mechanism.”

Lipid catabolism is regulated by autophagy (Singh et al., 2009). Additionally, autophagy promotes cancer survival by recycling intracellular proteins and organelles under conditions like oxidative stress or deprivation of nutrients (Tan et al., 2017; White, 2012), thereby sustaining overall metabolic homeostasis (Mizushima and Komatsu, 2011). Here we showed that shift toward lipid catabolism, under inhibited glutaminolysis, observed in our study, was accompanied by increased oxidative stress. We further confirmed that autophagy is triggered by the C.968 treatment. Given that autophagy is supporting fatty acid oxidation in the mitochondria and is considered crucial for the use of fatty acids delivered from adipocytes (Guo et al., 2013; Wen et al., 2017), our observation of C.968 treatment-induced oxidative stress (Figure 6A) and accelerated lipid catabolism (Figure 5) suggests that activation of the autophagy pathway was triggered to protect the cells from oxidative stress and to support lipid catabolism.

We showed that in the case of C.968 treatment, inhibition of one metabolic pathway is not sufficient to induce cancer cell death but rather leads to the activation of alternative pathways that facilitate cancer cell survival. By inhibiting glutaminolysis and autophagy, we blocked the induced “survival mechanism,” which resulted in significantly decreased cell viability. A recent study on pancreatic ductal adenocarcinoma and colon cancer also showed that glutamate shortage activates autophagy, and simultaneous inhibition of those pathways triggered apoptosis in that setting (Li et al., 2017; Seo et al., 2016), which corroborates our findings. Moreover, another recent studies showed that inhibition of either autophagy or fatty acid oxidation can block tumor recurrence (Viale et al., 2014), which further suggests that a combination of therapies that simultaneously target glutaminolysis and autophagy may be beneficial.

Taken together, our study provides a framework of how a metabolome-wide profiling approach can be used to identify metabolic adjustment of cancer cells to inhibited glutaminolysis and demonstrated a rational approach to identify and target molecular pathways that contribute to the activation of cancer survival mechanisms. This work provides fundamental insights into the adaptation of cancer cell metabolism to situations where glutamine metabolism is restricted and thereby reveals novel therapeutic targets that could be used in cancer treatment.

## Experimental Procedures

### Culture Conditions

The established cancer cell line MDA-MB-231, an in vitro model for TNBC, was obtained from the American Type Culture Collection (ATCC, Manassas, VA, USA). The cells were grown in Roswell Park Memorial Institute medium (RPMI-1640) (Sigma) supplemented with 10% fetal bovine serum, 1% penicillin–streptomycin solution (Sigma).

### Experimental Design

All experiments for the metabolomics study were performed in a 6-well plate format, and cells from two wells were merged to obtain the appropriate amount of material per single sample. The metabolomics study was performed in duplicate (for each duplicate, cells were harvested from two wells) in three independent experiments (in total six replicates were collected per condition). Cells were seeded in the 6-well plate at a density of 0.48 × 10^6^ cell per well. The medium was changed 24 h after seeding with a fresh 4 mL of conditioned media as follows: 1) fresh RPMI medium for control cells; 2) RPMI containing 0.05% of DMSO (vehicle) for control for treatment; 3) RPMI containing 5 μM concentration of C.968; and 4) RPMI containing 10 μM concentration of C.968. The cells were collected at 10 h, 24 h, 48 h, and 72 h after treatment. The cells used for counting and western blot were collected by trypsinization, and the harvesting and processing of cells for metabolite measurement are described below.

### Collection and Sample Processing for Metabolomics

The sample collection was performed as previously described with some modifications (Halama et al., 2013). Briefly, the medium was aspirated, and cells were washed twice with 37°C phosphate-buffered saline (PBS). After PBS was aspirated, 0.5 mL of 80% methanol in H_2_O was added per well, and the cells were scraped off of the well to simultaneously quench cellular processes and extract the metabolites. The scraped-in-methanol cells from two wells were combined in one Eppendorf tube (total volume of 1 mL), flash-frozen in liquid nitrogen, and then stored at −80°C until processing.

The metabolite extraction from the cells was performed in a series of freeze–thaw cycles. All samples were processed simultaneously. The samples were thawed on ice for 5 min with frequent vortexing and afterwards placed in liquid nitrogen for another 5 min. In total, samples underwent a series of three freeze–thaw cycles. After the final freeze–thaw cycle, the cells were centrifuged for 5 min at 18,000 ×g at 4°C. The supernatant was collected into the new tubes, frozen at −80°C, and shipped to Metabolon Inc. (Durham, NC, USA) on dry ice for metabolite measurements.

To account for differences in cell growth patterns caused by the treatment, the protein content from the pellet that remained after metabolic extraction was determined and used for normalization of the metabolic data. Sample processing for determination of the protein content was performed as previously described (Hansler et al., 2016). Briefly, pellets were dried in a speed vacuum for 20 min, and 100 μL of 0.2 M NaOH was added to each sample. Samples were heated for 20 min at 95°C with frequent vortexing and afterward centrifuged at 18,000 ×g for 5 min to remove debris. The protein content was determined using the Bio-Rad DC protein assay, relative to bovine serum albumin standards (0–1.2 mg/mL).

### Non-targeted Metabolic Profiling

Metabolic profiling was performed using Metabolon platforms deploying Waters ACQUITY ultra-performance liquid chromatography (UPLC) and a Thermo Scientific Q-Exactive high-resolution/accurate mass spectrometer interfaced with a heated electrospray ionization (HESI-II) source and Orbitrap mass analyzer, as previously described (Evans et al., 2009).

Briefly, recovery standards were added for quality control purposes, and the extract was divided into the following fractions: 1) two fractions for analysis by two separate reverse-phase (RP)/UPLC-mass spectrometry (MS)/MS methods with positive ion mode electrospray ionization (ESI); 2) one fraction for analysis by RP/UPLC-MS/MS with negative ion mode ESI; 3) one fraction for analysis by hydrophilic interaction chromatography (HILIC)/UPLC-MS/MS with negative ion mode ESI; and 4) one fraction reserved for backup.

The sample extract was dried under nitrogen and reconstituted in solvents compatible with each of the four methods. The aliquots were analyzed in the following conditions: 1) acidic positive ion (optimized for hydrophilic compounds)—extract gradient eluted from a C18 column (Waters UPLC BEH C18-2.1 × 100 mm, 1.7 μm) with water and methanol containing 0.05% perfluoropentanoic acid and 0.1% formic acid; 2) acidic positive ion (optimized for hydrophobic compounds)—extract gradient eluted from C18 (Waters UPLC BEH C18-2.1 × 100 mm, 1.7 μm) with methanol, acetonitrile, water, 0.05% perfluoropentanoic acid, and 0.01% formic acid; 3) basic negative ion—extract gradient eluted from a separate dedicated C18 column using methanol and water containing 6.5 mM ammonium bicarbonate at pH 8; and 4) negative ionization—extract gradient eluted from a HILIC column (Waters UPLC BEH Amide 2.1 × 150 mm, 1.7 μm) using water and acetonitrile with 10 mM ammonium formate at pH 10.8. In the MS analysis, the scan range varied between methods but covered the range of 70–1000 m/z.

Metabolon’s hardware and software were deployed for raw data extraction, and compounds were identified by comparison of peaks to library entries of purified standards based on retention index, accurate mass match to the library ± 10 ppm, and MS/MS forward and reverse scores between the experimental data and authentic standards. Library matches for each compound were checked for each sample. The metabolic data was normalized to correct variations resulting from inter-day tuning differences in the instrument. Each compound was corrected in a run-day.

### Cell Respiration Assay

The OCR was measured to monitor whether (1) C.968 inhibited glutaminolysis in MDA-MB-231 and (2) fatty acids served as an energy source for MDA-MB-231, using the XF96 extracellular flux analyzer (Seahorse, Agilent). The assay was prepared according to the Seahorse protocols.

To monitor the inhibition of GAC by C.968, the cells were seeded one day before the experiment in the 96-well XF Cell Culture Microplate (Seahorse, Agilent) at a cell density of 15,000 cells/well in RPMI-1640 media (Sigma). The medium was changed, and the cells were treated with vehicle or 10 μM C.968 for 10 h. One hour before the measurements, medium was changed to base medium (Seahorse, Agilent) supplemented with: (1) glucose (10 mM); (2) glutamine (2 mM); or (3) a mixture of glucose and glutamine (10 mM and 2 mM) adjusted to pH 7.4, and the cells were placed in the non-CO_2_ incubator for 1 h. The OCR was measured over a period of 80 min after sequential injection of oligomycin (1 μM), FCCP (1 μM), and antimycin A/rotenone (0.5 μM), at time points provided in the manual.

To monitor cell ability to use fatty acids, the cells were seeded in 96-well XF Cell Culture Microplate (Seahorse, Agilent) at a cell density of 15,000 cells/well in growth medium RPMI-1640 (Sigma) and cultivated for 24 h. The medium was replaced with a base medium (Seahorse, Agilent) supplemented with 0.5 mM glucose, 1 mM GlutaMAX, 0.5 mM carnitine, and 1% fetal bovine serum, and the cells were cultivated for 24 h. At 1 h before the measurements, cells were washed once with Krebs–Henseleit buffer (111 mM NaCl, 4.7 mM KCl, 1.25 mM CaCl_2_, 2 mM MgSO_4_, 1.2 mM NaH_2_PO_4_, 2.5 mM) supplemented with glucose, 0.5 mM carnitine, and 5 mM HEPES and incubated in the same media for 45 min in the non-CO_2_ incubator. At 20 min prior to the measurements, etomoxir (40 μM) or DMSO was added to the wells, and cells were incubated for next 15 min. Right before taking the measurements, XF Palmitate-BSA FAO substrate was added to the wells. The oxygen consumption rate was measured over a period of 80 min after sequential injection of oligomycin (1 μM), FCCP (1 μM), and antimycin A/rotenone (0.5 μM), at the time points provided in the manual.

### Western Blotting

Whole cell lysates were prepared in a 20 mM Tris-HCL lysis buffer (pH 7.49) supplemented with protease–phosphatase cocktail inhibitors (Roche) and phenyl methane sulfonyl fluoride (Sigma Aldrich). Samples were lysed by three freeze–thaw cycles in liquid nitrogen as previously described (Halama et al., 2016). The supernatant was collected after centrifugation for 10 min, and the total protein content was quantified using the DC protein assay (Bio-Rad). The proteins were denatured by incubation with 4× Laemmli buffer at 95°C for 10 min.

Gel electrophoresis was performed using the prepared whole cell lysate and transferred to a polyvinylidene fluoride membrane (Bio-Rad). The membrane was blocked in 5% milk solution in Tween-PBS (PBS with 0.1% Tween 20) for one hour and incubated overnight in a primary antibody at 4°C. The overnight incubation was followed by three washing steps in Tween-PBS. The membrane was incubated in the corresponding secondary antibody followed by three washing steps. Both primary and secondary antibodies were prepared in the recommended dilution in a 5% milk solution in Tween-PBS. The blots were developed and visualized under a ChemiDoc system (Amersham, Bio-Rad, USA). The primary antibodies used were CPT1a (Cell Signaling, #12252S) and CPT2 (Santa Cruz, #sc-377294). The corresponding secondary antibodies included horseradish peroxidase–conjugated anti-mouse (Cell Signaling) and anti-rabbit (Cell Signaling).

### Cell Cycle Monitoring

MDA-MB-231 cells were exposed to two different concentrations of C.968 (5 μM and 10 μM) for 10 h, 24 h, 48 h, and 72 h. Cells were washed, fixed with 70% ethanol, and incubated for 30 min at 37°C with 0.1 % RNase A in PBS. Cells were washed again, resuspended, and stained with 25 μg/mL PI for 30 min in PBS at room temperature. The cell distribution across the cell cycle was analyzed by flow cytometry using a BD LSRFortessa analyzer (BD Biosciences) as previously described (Siveen et al., 2014).

### Cell Death Monitoring

MDA-MB-231 cells were exposed to two different concentrations of C.968 (5 μM and 10 μM) for 10 h, 24 h, 48 h, and 72 h. The cells were collected by trypsinization, washed with PBS, and stained with fluorescein-conjugated annexin V and PI (BD Biosciences). The percentage of cells undergoing apoptosis or necrosis was measured by flow cytometry using a BD LSRFortessa analyzer (BD Biosciences) as previously described (Hussain et al., 2007). The cell viability status was quantified based on the staining as follows: the live (Annexin-FITCnegative, PI-negative), early apoptosis (Annexin-FITC^+^positive, PI-negative), late apoptosis (Annexin-FITC^+^ positive, PI^+^ positive) apoptotic cells, and necrosis (Annexin-FITC negative, PI^+^positive).

### Statistical Analysis

Statistical analysis was performed in R version 3.1.3 and R-Studio version 0.97.551. The metabolomics data were normalized to protein content, followed by log transformation and z-scoring.

Multivariate linear regression was used to assess the statistical significance of the association of metabolites with C.968 treatment in MDA-MB-231 over the entire course of the experiment while adjusting for time. Additionally, we used linear regression to test the association of metabolites with the C.968 treatment in MDA-MB-231 cells at specific time points separately (10 h, 24 h, 48 h, and 72 h). Significance was defined at a stringent Bonferroni level of p⍰<⍰1.44⍰×⍰10^−04^ (p⍰<⍰0.05/346).

We also performed analysis of ratios between metabolites because using ratios reduces the overall variance and allows for identification of biologically relevant pathways, as previously reported (Altmaier et al., 2008; Petersen et al., 2012). We examined the time-resolved effect of the C.968 treatment on metabolite ratios in MDA-MB-231 cells, using a multivariate linear regression model for each pair of metabolites. In total, we calculated 68,635 metabolic ratios (346⍰× 345/2). We considered metabolite ratios as significantly altered if they satisfied both the following conditions: p-value⍰<⍰8.37⍰×⍰10^−07^(p⍰<⍰0.05/59,685) and p-gain⍰(define measure to identify statistically significant metabolite ratios (Petersen et al., 2012)) >⍰5.96⍰×⍰10^05^ (p-gain⍰>⍰10⍰×59,685).

The Orthogonal Projections to Latent Structures (OPLS) analysis was performed with SIMCA version 14 (Umetrics, Umea, Sweden).

## Acknowledgments

This study was supported by QNRF grant NPRP8-061-3-011 and also by ‘Biomedical Research Program’ funds at Weill Cornell Medicine in Qatar, a program funded by the Qatar Foundation. The funders had no role in the study design, data collection and analysis, decision to publish, or preparation of the manuscript. We thank Ms. Aleksandra M. Liberska (flow cytometry supervisor) and the Flow Cytometry Facility within the Microscopy Core at Weill Cornell Medicine-Qatar for contributing to these studies.

## Competing Interests

The authors declare no competing financial interests.

## Author Contributions

Conceptualization, K.S., A.H., G.K., O.E., S.S.G., and S.U.; Investigation, A.H., M.K.; Methodology, A.H., M.K., S.B.Z., S.S.D., K.S.S., A.I., and N.J.S.; Writing and Revisions, A.H., K.S., M.K., S.U., S.B.Z, S.S.G., A.M.B, G.K., and O.E.; Funding Acquisition, K.S.; Supervision, K.S., S.S.G., G.K., and O.E.

---

**Supplementary Table 1.** Metabolite information.

**Supplementary Table 2.** DMSO has minor effects on cancer cell metabolism. Differences between control and vehicle treated cells were tested for significance at 10h, 24h, 48h and 72h. All measured metabolites together with FDR and Bonferroni corrected p-values were included. Supplementary Figure 1. **A)** Western blot analysis of the glutaminase C (GAC). **B)** Phase-contrast microscopy of MDA-MB231 cells at 72h after treatment with C.968.

**Supplementary Figure 2.** DAPI staining of MDA-MB231 cells at 10h, 24h, 48h and 72h after treatment with C.968 (5μM and 10 μM).

**Supplementary Figure 3.** The box plots present alteration patterns of metabolites involved in lipid metabolism after treatment of cells with C.968. Vehicle treated cells are depicted in grey; treated with 5μM in green and treated with 10 μM C.968 in dark green. Bonferroni significant (p-value ≤ 1.35x10^−4^) changes are highlighted on the green background (for decrease) and on the red background (for increase). The alterations showing changes which are not Bonferroni significant are highlighted on the dotted green (for decrease) and dotted red (for increase) background respectively.

